# Artificial physics engine for real-time inverse dynamics of arm and hand movement

**DOI:** 10.1101/2023.02.07.527431

**Authors:** Mykhailo Manukian, Serhii Bahdasariants, Sergiy Yakovenko

**Affiliations:** Faculty of Applied Science, Ukrainian Catholic University, Lviv, Ukraine; Department of Human Performance—Pathophysiology, Rehabilitation, and Performance, School of Medicine, West Virginia University, Morgantown, West Virginia, United States; Department of Neuroscience, School of Medicine, West Virginia University, Morgantown, West Virginia, United States; Rockefeller Neuroscience Institute, School of Medicine, West Virginia University, Morgantown, West Virginia, United States; Mechanical and Aerospace Engineering, Benjamin M. Statler College of Engineering and Mineral Resources, West Virginia University, Morgantown, West Virginia, United States; Department of Biomedical Engineering, Benjamin M. Statler College of Engineering and Mineral Resources, West Virginia University, Morgantown, West Virginia, United States

## Abstract

Simulating human body dynamics requires detailed and accurate mathematical models. When solved inversely, these models provide a comprehensive description of force generation that evaluates subject morphology and can be applied to control real-time assistive technology, for example, orthosis or muscle/nerve stimulation. Yet, model complexity hinders the speed of its computations and may require approximations as a mitigation strategy. Here, we use machine learning algorithms to provide a method for accurate physics simulations and subject-specific parameterization. Several types of artificial neural networks (ANNs) with varied architecture were tasked to generate the inverse dynamic transformation of realistic arm and hand movement (23 degrees of freedom). Using a physical model to generate the training and testing sets for the limb workspace, we developed the ANN transformations with low torque errors (less than 0.1 Nm). Multiple ANN implementations using kinematic sequences solved accurately and robustly the high-dimensional kinematic Jacobian and inverse dynamics of arm and hand. These results provide further support for the use of ANN architectures that use temporal trajectories of time-delayed values to make accurate predictions of limb dynamics.

## Introduction

The accurate and fast models of body dynamics help to unravel neuromechanical interactions within movement control pathways and to develop intuitive human-machine interfaces. Yet, the simulation speed–accuracy tradeoff is challenged by the body’s complex segmental dynamics and the quality of subject morphometry. In the context of the hand and arm, a realistic musculoskeletal model consists of about 23 degrees of freedom (DOF) actuated by 52 force-generating musculotendon units scaled by limb segment geometry. This complexity is a challenge for real-time simulations, for example, for brain-computer interfaces (BCI), and can benefit from approximations of musculoskeletal transformations (Sobinov et al. 2020; Smirnov et al. 2021) and limb physics. Artificial neural networks (ANNs) were previously applied to solve this problem by approximating the input-output transformations from neural signals to the intended movement (Hochberg et al. 2006; Donoghue et al. 2007; Collinger et al. 2013; Wodlinger et al. 2015). However, these ANN-based solutions have not exceeded the robust decoding of several (up to 10) DOFs even within limited, stereotypical tasks and postures (Wodlinger et al. 2015). The potential reason for the limited success is the expectation that the neural-to-motion transformation can be solved *without* the explicit description of segment dynamics and interactions with external objects. For example, in human BCIs, optimized decoders trained only on the kinematics of reaching to objects are challenged by the presentation of physical objects ^9,109,10^(Downey et al. 2017). The neural activity likely expresses nonlinearities of muscle actions and their dependency on dynamical demands of object manipulation, which would disrupt simplistic statistical decoders. Theoretically, increasing decoder complexity and training with large datasets can mitigate this problem. However, the exponential increase in both size of the training dataset and the required training duration may be prohibitive for realistic transformations. While ANN formulation is attractive for developing neural interfaces for movement and motor learning, it has fallen short of capturing motor complexity.

In this study, we continue the effort of developing a theoretical framework that employs ANNs guided by signal processing in BCI to define a full transformation from neural inputs to decoded physical actions. The main contribution of this work is the methodology for separately solving spatiotemporal dynamics corresponding to limb physics that can be combined with our previous solutions of musculoskeletal relationships (Sobinov et al. 2020; Smirnov et al. 2021). The ANN-based solutions of spatiotemporal characteristics in dynamical systems have been recently shown for the relatively simple Lorenz system (Park et al. 2022). The representation of accurate arm dynamics with the ANN formulation is the focus of this study. Preliminary results of this work were published in the thesis form (Manukian 2021).

## Materials & Methods

### Kinematic and kinetic datasets

Joint kinetics and kinematics representing movements within the arm workspace were simulated and used for training and testing of the model. Similar to (Graziano 2006), the workspace was explored with a set of movements. All combinatorial combinations of limb movements within a 3×3×3 grid (Fig.1) were simulated with a bell-shaped velocity profile imposed on the straight end-point trajectory (Kang, He, and Tillery 2005) for three durations (0.5, 1.0, 2.0 s). The initial and final joint postures for each vertex in the grid were based on a set of representative observations. The vertices were selected as follows: middle-right vertex corresponded to the arm outstretched forward and parallel to the ground (posture in Fig. 1A); top-right and bottom-right vertices corresponded to 45º vertical deviations from this position; top-, middle-, and bottom-left vertices tracked the maximum outstretched arm at the same level as on the right and aligned with the left shoulder. The proximal vertices were chosen at the same vertical level as the distal vertices but placed 10 cm from the body. The middle positions were chosen midway between the proximal and distal vertices. The set of joint kinetics was computed at 1 ms timestep using a custom 23 DOF biomechanical model of the human arm and hand (Fig. 1B), formulated in Simscape Multibody (Simulink R2022a, MathWorks) and scaled to a representative subject morphometry (Winter 2009, 86). The end-point trajectories were simulated with shoulder flexion, shoulder internal-external rotation, and elbow flexion; other joints remained in a neutral posture (the middle of the range of motion). The resulting N=351 reaching movements were organized as the data structures containing [*q,q’,q’’*,τ], where *q* is the vector of joint angles, *q’* and *q’’* are its derivatives, and τ is the joint torque. Velocity and acceleration values of not moving degrees of freedom (zero values) were excluded from the state vector of 69 kinematic and 23 kinetic values.

**Fig. 1:**
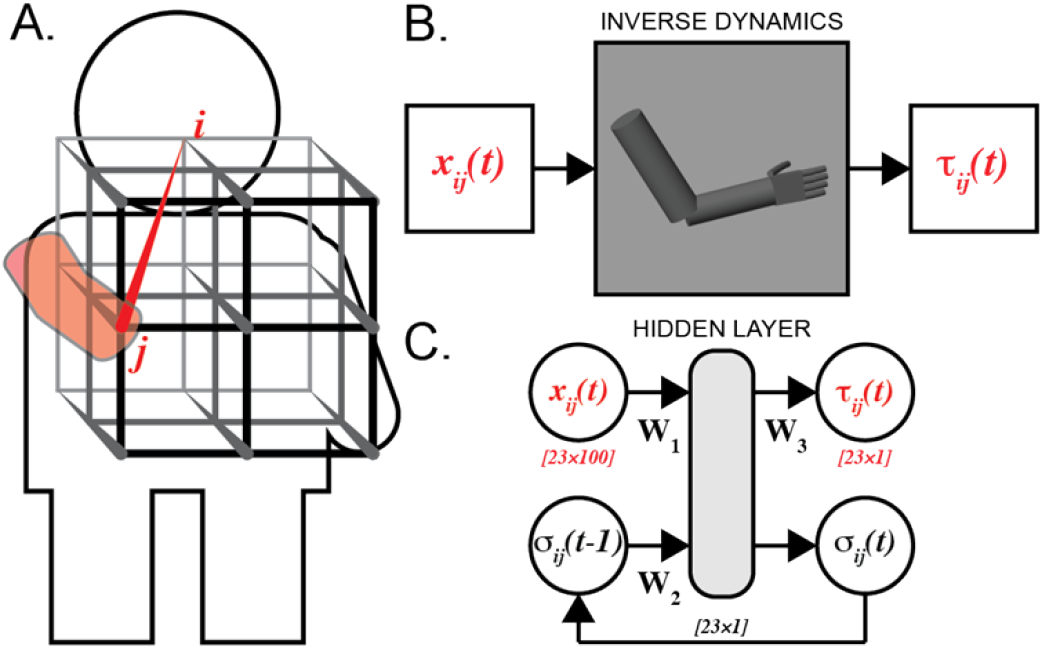
Dataset based on the movement within a 3D grid (A) was computed with an inverse dynamic model of segmented arm and hand (B). The recurrent neural networks (RNNs) (C) were used to approximate the transformation: *x*_*ij*_ and τ_*ij*_ are input kinematic and output kinetic signals, respectively; σ_*ij*_ is output of the hidden layer fed back to compute next-step kinetics; **W**_1-3_ are weight matrices.

### Training, validation, and testing subsets

A total of 1,124,000 state vectors, representing all possible movement combinations, were split into training (71%), validation (15%), and testing (14%) subsets. We used one input sequence length (100 samples, fetched from state vectors) to train ANNs and additional three lengths (10, 20, 50 samples) to examine the relative performance of transformations. The accuracy of the ANNs during training process was monitored with validation subset. The testing subset was unseen by ANN during the training and was used post-hoc for the same purposes. Data representing different movement durations were equally distributed among the subsets. The z-score normalization was applied to input torque values to ensure their scaled contribution to ANN predictions. The predicted torques were then de-normalized to evaluate absolute errors across modeled DOFs.

### Hyperparameters

A preliminary search with a grid pattern was performed over four hyperparameters: learning rate (0.0001, 0.0005, or 0.001), the hidden layer size (23, 69, or 115), the number of recurrent layers (1, 3, or 5), and the network architecture type. The search resulted in 54 experiments that were limited to training with only 15 epochs (each epoch is the use of a full training dataset). We considered the following three ANN types: Elman RNN (Elman 1990), long short-term memory cell (LSTM) (Sundermeyer, Schlüter, and Ney 2012), and gated recurrent unit (GRU) (Dey and Salem 2017), excelling at statistical forecasting (Hewamalage, Bergmeir, and Bandara 2021).

### Statistical Analysis

The mean squared error (MSE) between target and predicted torques was used to monitor ANN training. The final accuracy was measured with root MSE (RMSE). The normality was tested with a one-sample Kolmogorov-Smirnov test, and Kruskal-Wallis test was used to evaluate differences in non-normally distributed groups. The ANN execution time was measured on CPU and GPU using the Google Colab Pro environment. The ANN resistance to numerical noise was performed by perturbing test data with Gaussian noise. All analyses were conducted using PyTorch (Paszke et al. 2019) and SciPy 1.0 (Virtanen et al. 2020) software with the significance level (α) set to 5%.

## Results

### Hyperparameter Tunning

Effective training and accurate approximations require optimization of many hyperparameters. To focus on the appropriate domain, we performed a preliminary hyperparameter grid search (see *Hyperparameters* in Methods). We found that the most critical hyperparameter was the learning rate. Consistent with the previous work (Wilson and Martinez 2001), the use of slower values below 5×10^−4^ was less optimal–decreasing accuracy gain per epoch and increasing the training time. The learning rates over 5×10^−4^ resulted in less accurate and less dynamically stable solutions. The next parameter addressed in preliminary optimization was the number of hidden layers. There was no benefit to the deep structure of ANN, which resulted in the exclusion of the most three- and five-layer configurations that suffered from the overfitting and high inaccuracies in testing conditions.

### Approximating Inverse Dynamics

Five candidate ANNs approximated solutions to the equations of motion describing the reaching movements in the 23 DOF arm and hand model. For all networks, the optimal learning rate was equal to 5×10^−4^. Fig. 2 shows the best examples of a single-trial simulations with the lowest errors. Torques of proximal and distal joints show a close relationship with the reference trajectories (dashed). Contrasting these results to the worst performance shown in Fig. 3, not all ANNs performed equally. The torque trajectories computed with the Elman RNN were prone to oscillatory behavior, as seen in Fig. 3 A, C, and F. Similarly, the single layer LSTM could generate large transient deviations, as seen in Fig. 3A. Both Elman RNN and LSTM transformations were prone to amplitude scaling errors. The synthesis of performance across all trials in the test dataset is reported in Table 1. The best accuracy was attributed to the one-layer GRU with 115 computational nodes compared to other tested architectures (p<5×10^−6^, Fig. 4). Elman RNN expectedly performed least accurately as compared to other ANNs for 100 sample sequences (p<7×10^−21^).

**Table 1.** The relationship of input length with accuracy for different ANN types. The most accurate candidates are marked in **bold**.

| ANN Architecture | Torque RMSE, Nm |  |  |  |
| --- | --- | --- | --- | --- |
|  | 10 samples | 20 samples | 50 samples | 100 samples |
| RNN-1-115 | 0.2870 | 0.1180 | 0.0757 | 0.0736 |
| GRU-1-69 | 0.4515 | 0.3094 | 0.1149 | 0.0433 |
| LSTM-3-69 | 1.1822 | 0.4339 | 0.0780 | 0.0398 |
| GRU-1-115 | 0.3503 | 0.1837 | 0.0633 | **0.0273** |
| LSTM-1-115 | **0.2764** | **0.0872** | **0.0401** | 0.0312 |

**Fig. 2:**
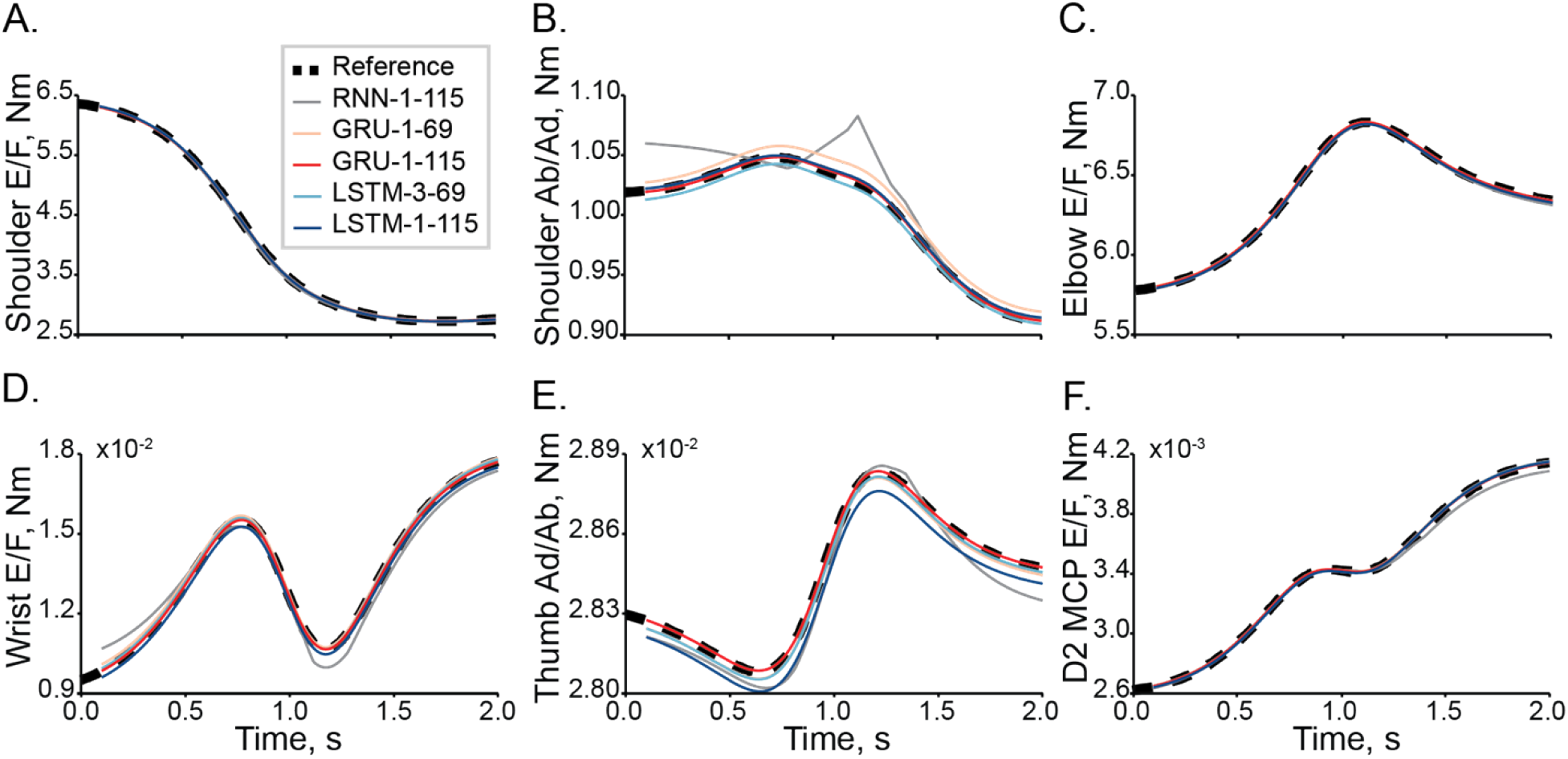
The best example with the lowest errors of inverse dynamic transformation in a single trial. Joint torques are shown for a selection of DOFs: (A) shoulder flexion, (B) shoulder adduction, (C) elbow flexion, (D) wrist flexion, (E) thumb abduction, and (F) index finger flexion. Used naming convention is XXX-Y-ZZZ, where XXX is ANN type, Y is the number of hidden layers, and ZZZ is the number of the computational nodes within one hidden layer; type “RNN” refers to Elman RNN.

**Fig. 3:**
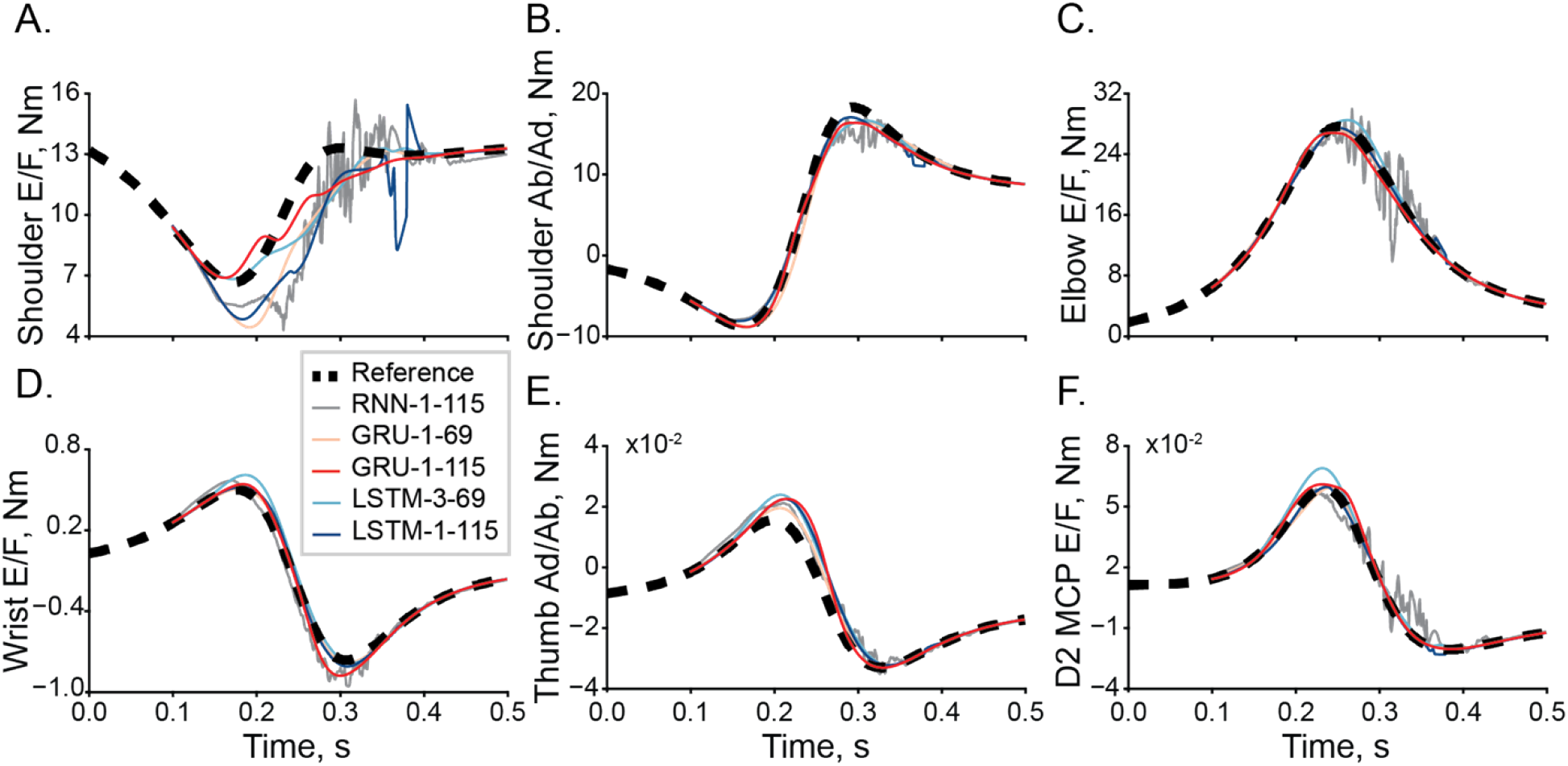
The worst example with the highest errors of inverse dynamic transformation in a single trial. The organization is the same as in Fig.2.

**Fig. 4:**
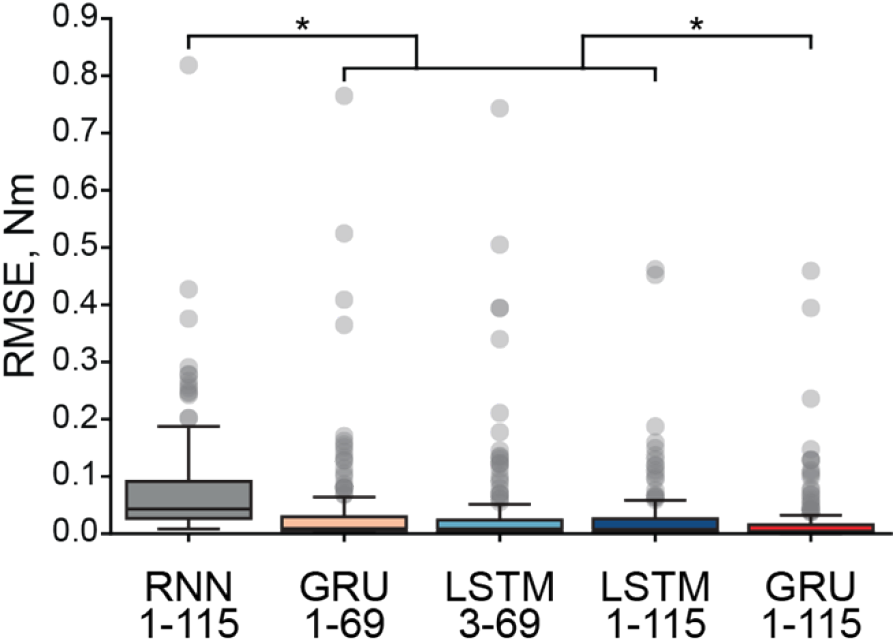
Simulated torque errors with different ANN architectures. The box plots show the 25th and 75th percentiles of the RMSE distributions between target and simulated torques, solid horizontal lines are medians; whiskers indicate full range, and circles mark outliers (1%). The significant differences tested with the Mann-Whitney pairwise comparison (α < 5%) are indicated with *.

### Input Sequence Length

We used one input sequence length (100 samples) to train ANNs and tested them with four lengths (10, 20, 50, and 100) to examine the relative performance of transformations. The accuracy increased with the increase of sequence length, as shown by the mean predicted torque RMSE values in Table 1. This trend saturated for sequences of 50 samples and longer, resulting in <0.1 Nm errors for all ANNs. At shorter lengths (*n*=20), RNN transformation outperformed GRU and the three-layer LSTM transformations. However, the performance did not improve as quickly as with other ANN types with longer sequences. Overall, both one-layer LSTM and GRU with 115 nodes delivered the highest accuracy.

### Computation Latency

We measured the transformation latency per one output sample required by each ANN type to evaluate the real-time readiness of the current implementation using CPU and GPU hardware. On GPU, one-layer ANNs processing up to 50 sample sequences were executed faster than real-time (<1 ms per sample). For implementations with 100 sample sequences, the latency declined to just under 2 ms per sample. The three-layer LSTM was faster than real-time only in the 10-sample mode and took about 5.5 ms per sample in the 100-sample mode. On CPU, one-layer ANNs executed in real-time only with 10 sample sequences and not any longer sequences. For the mode with 100 samples, the latencies were about 6 ms per sample, and the RNN was the fastest executing at 2 ms per sample. The three-layer architecture was slow in 10 (∼2 ms) and 100 (∼14 ms) sample modes.

### Numerical Noise

We examined ANN tolerance to noise in models with 100 sample input sequences by adding 1% and 5% of peak-to-peak Gaussian noise to kinematic trajectories (Table 2). The accuracy of Elman RNN decreased 1.34x with 1% noise and 3.73x with 5% noise relative to simulations without noise. The accuracy decreases were 1.2x and 3.07x for the 69-node GRU and 1.31x and 3.81x for the 115-node GRU. The absolute error values, however, were lower in GRU 1-115 than those in GRU 1-69 with noise. The LSTM transformations were the most sensitive to noise, with decreases of 2x and 6.81x for the single-layer structure and decreases of 8.55x and 21.05x for the triple-layer structure.

**Table 2.** The relationship between input noise and accuracy for different ANN types.

| ANN Architecture | Torque RMSE, Nm |  |
| --- | --- | --- |
|  | 1% Noise | 5% Noise |
| RNN-1-115 | 0.1030 | 0.2987 |
| GRU-1-69 | 0.0616 | 0.1827 |
| LSTM-3-69 | 0.4770 | 1.2050 |
| GRU-1-115 | **0.0438** | **0.1433** |
| LSTM-1-115 | 0.0775 | 0.2765 |

## Discussion

In this study, we developed and tested multiple ANN architectures to solve the spatiotemporal inverse dynamics of a segmented arm and hand. The numerical model of arm dynamics was used to create input-output datasets for the physiological workspace of arm movements. The trained ANNs were examined for the accuracy, robustness, and execution efficiency to validate ANN approach and to examine the impact of different parameters. We demonstrate that the single-layer LSTM and GRU models with 115 nodes can perform accurate and real-time simulations. These simulations executed on cloud service GPU with 50 ms sequences produce the most accurate results. The same models can run in real-time on cloud service CPU with short sequences (10 samples) and, consequently, less accuracy. The additional testing with noisy inputs demonstrated robustness of models to resist noise generating less than 0.05 Nm errors with the additional 1% noise that increased three-fold with the additional 5% noise (see Table 2, GRU model).

### Optimal Network Structure

We have conducted a preliminary parametric search of ANN structures simulating high dimensional (23 DOF) inverse dynamics of human arm and hand. This limited our analysis to five candidates that were expected to generate *“good-enough”* approximations. Fig. 2 and 3 show the performance of the ANNs. While the performance across all ANNs was surprisingly accurate, the Elman RNN model generated oscillatory predictions suggesting that the training might have reached a suboptimal solution near a local minimum, for example, near saddle type critical points. With the increasing realism in biomechanics of human limbs, these saddle points proliferate, presenting a challenge to ANN training (Pascanu et al. 2014). The methods to overcome it exist (Dauphin et al. 2014); however, they have not been implemented in modern programming frameworks. A second issue with the Elman RNN model was the saturation of performance when the training MSE approached zero. This problem is termed the “vanishing gradients” problem, apparent in the relatively high errors in Fig. 4. GRU and LSTM model types were not prone to this methodological limitation. In LSTM models, scaling errors were minimal; however, the outcomes had infrequent dynamical transients (e.g., Fig. 3A), which would be destabilizing in forward dynamic simulations. These transients indicate the overfitting that occurs in LSTMs with many computational nodes (Lawrence, Giles, and Tsoi 1997). The LSTM with fewer nodes did not suffer from this problem in support of this idea. The GRU model outputs were smooth with only minor scaling and dynamical errors. This high performance is likely due to the GRU algorithmic complexity, which lies between those of Elman RNN and LSTM demonstrating diminishing returns of both simple and complex architectures in this problem and demonstrates minimized probability of overfitting (Gruber and Jockisch 2020). The one hidden layer GRU was the most accurate with LSTM as a close second. Therefore, GRU and LSTM topologies are best suited for solving inverse dynamics for realistic limb biomechanics.

### Sequences for neural net training

Many behaviors are dynamical, history-dependent, and produced by sequences of actions. All goal-directed movements are a result of sequential and coordinated actions of the musculoskeletal system. For example, the actions of walking and reaching to a target are generated by the spatiotemporal sequences of cortical ensembles and muscle groups generating forces that result in the progression of mechanical actions (Yakovenko and Drew 2015). The planning and execution within neural control pathways are thought to require both inverse and forward computations to generate appropriate control sequences and to monitor their execution through the comparison of expected and sensed signals. This general embedding of control structure within neural pathways is supported by the theory of internal models (Wolpert, Ghahramani, and Jordan 1995; Gribble and Scott 2002). Historically, the ANN approach was envisioned as the simplified formulation of neural computations (Richards et al. 2019), but the current implementations are limited compared to neural functions (Zador et al. 2022). For ANNs to solve the control of movement, the essential computations need to solve musculoskeletal limb kinematics and history-dependent limb dynamics.

Our result demonstrates that the use of sequences in training and evaluation of ANN models for inverse dynamics is a valid approach. LSTM and GRU models for natural language processing (Mikolov and Zweig 2012; Mikolov et al. 2011) and kinematic video tracking (Gordon, Farhadi, and Fox 2018) embed history-dependent relationships within sequences and are less sensitive to the problem of “catastrophic forgetting” (Jedlicka et al. 2022). Similarly, the Newtonian relationship through second-order differential equations governing joint movement imposes the history-dependence on kinematic sequences. This study provides the first demonstration of realistic arm and hand dynamics with the use of ANNs with memory capabilities to represent sequences.

We extend the previous studies focused on relatively low-dimensional dynamics of industrial robotic systems to the realistic high-dimensional arm and hand movements. Previously, inverse dynamic transformations with ANNs were typically limited to several degrees of freedom for the control of robotic devices (e.g., Kuka arm robot, 5-7 DOFs) (Liu et al. 2019; Rueckert et al. 2017). Yet, we achieve similar torque prediction performance (less than 0.15 Nm errors with high level of noise) as in these low-dimensional systems (Çallar and Böttger 2022). Furthermore, we propose this approach as the extension for the full musculoskeletal dynamics that details moment arm and muscle length relationships with posture (Smirnov et al. 2021). This combination of physically explainable ANN models can, thus, provide decoding of motor intent or the control of wearable powered robotics using muscle-level resolution of biomechanical state.

### Limitations

We chose to perform a heuristic grid-search method to identify essential parameters and select them as hyperparameters for further testing. Automating the design of ANN can outperform custom architectures (Zoph et al. 2017; Feurer and Hutter 2019). For example, the optimization of hidden layers and nodes (Stathakis 2009) within the optimization of accuracy and decreased model complexity can find ANN structures that outperform the current result. Overall, the full parameter space is currently unexplored for the applications that describe limb dynamics. Another potential limitation is the limited exploration of behavioral space in this and other previous studies. While we have introduced a generalized approach to the representation of workspace for reaching movements. Other behavioral tasks may be challenging for this model. For example, grooming, object interactions, or pathologies like tremor (high-speed 8 Hz oscillations) may need to be included in the training dataset for specific applications. The movements with sharp direction reversals may require models with shorter than 50 ms (Table 2) input sequences. We showed that a single-layer LSTM could maintain prediction accuracy (about 0.1 Nm error) with only 20 ms input sequences indicating that the proposed method remains valid with the outlined mitigation strategies.

## Conclusions

In this study, we trained and optimized a machine learning algorithm to approximate solutions to the complex inverse dynamic problem. The ANNs were tested on a 23-DOF human arm and hand model. We demonstrated that simple one-layer ANNs with memory is real-time accurate and noise-tolerant when predicting kinetics in complex multi-joint systems. Together, these findings suggest an instrumental role for ANNs in the development of control algorithms that approximate dynamic simulations with minimal latencies, thus enabling accurate and computationally inexpensive control of wearable prosthetics and assistive devices.

## Acknowledgements

This work was supported by the Office of the Assistant Secretary of Defense for Health Affairs through the Restoring Warfighters with Neuromusculoskeletal Injuries Research Program (RESTORE) under Award No. W81XWH-21-1-0138.

